# Vps54 regulates *Drosophila* neuromuscular junction development and controls postsynaptic density composition via a Rab7-dependent mechanism

**DOI:** 10.1101/2020.02.17.952721

**Authors:** Prajal H. Patel, Emily C. Wilkinson, Emily L. Starke, Malea R. McGimsey, J. Todd Blankenship, Scott A. Barbee

**Author notes:** These authors contributed equally (Prajal H. Patel, Emily C. Wilkinson).

## Abstract

Vps54 is a subunit of the Golgi-associated retrograde protein (GARP) complex, which is involved in tethering endosome-derived vesicles to the *trans*-Golgi network (TGN). In the wobbler mouse, a model for human motor neuron (MN) disease, reduction in the levels of Vps54 causes neurodegeneration. However, it is unclear how disruption of GARP-mediated vesicle transport leads to MN dysfunction and ultimately neurodegeneration. To better understand the role of Vps54 in MNs, we have disrupted expression of the *Vps54* ortholog in *Drosophila* and examined the impact on the larval neuromuscular junction (NMJ). Here, we show that both null mutants and MN-specific knockdown of *Vps54* leads to NMJ overgrowth. Reduction of *Vps54* partially disrupts localization of the t-SNARE, Syntaxin-16, to the TGN but has no impact on endosomal pools. Presynaptic knockdown of *Vps54* in MNs combined with overexpression of the small GTPases Rab5, Rab7, or Rab11 suppresses the *Vps54* NMJ phenotype. Conversely, knockdown of *Vps54* combined with overexpression of dominant negative Rab7 causes axonal and behavioral abnormalities including a decrease in postysynaptic Dlg and GluRIIB levels without any effect on GluRIIA. Taken together, these data suggest that *Vps54* controls larval MN axon development and postsynaptic density composition by modulating Rab7-mediated endosomal trafficking.

## INTRODUCTION

Endocytic trafficking is critical for many specialized processes in neurons including axon growth, guidance, plasticity, and for the maintenance of cellular homeostasis (Wojnacki and Galli, 2016). Disruption of endocytic trafficking pathways can cause neurodevelopmental defects and contribute directly to neurodegeneration (Schreij et al., 2016). A destabilizing point mutation in the gene encoding the vacuolar protein sorting-associated protein 54 (*Vps54*) is responsible for age-progressive motor neuron (MN) degeneration observed in the wobbler mouse model for human MN disease (Schmitt-John et al., 2005). *Vps54* loss-of-function in the mouse causes embryonic lethality and is characterized by the underdevelopment of MNs and cardiac muscle. Vps54 is a core subunit of the evolutionarily conserved Golgi-associated retrograde protein (GARP) complex, which localizes to the *trans*-Golgi network (TGN) and is involved in tethering retrograde transport carriers derived from endosomes (Bonifacino and Hierro, 2011). It has been postulated that the reduction of Vps54 levels, and thus a compensatory disruption of the GARP complex, contributes directly to MN degeneration (Moser et al., 2013; Schmitt-John, 2015).

GARP is a ubiquitously expressed complex composed of the Vps51, Vps52, Vps53, and Vps54 proteins (Conibear and Stevens, 2000). Within this complex, the Vps54 N-terminus binds to soluble *N-*ethylmaleimide-sensitive fusion protein attachment protein receptors (SNAREs). Vps54 interacts with SNAREs involved in retrograde transport including: Syntaxin-6 (Stx6), Syntaxin-16 (Stx16), and Vamp4 and is required for t-SNARE assembly (Perez-Victoria and Bonifacino, 2009). In contrast, the C-terminal domain interacts with endosomes and is dispensable for both GARP and t-SNARE complex formation (Quenneville et al., 2006). Knockdown of Vps54 and other GARP subunits result in defects in retrograde transport of vesicles including those carrying the mannose-6-phosphate receptor (M6PR) and some v-SNAREs (Conibear and Stevens, 2000; Perez-Victoria and Bonifacino, 2009; Perez-Victoria et al., 2008; Quenneville et al., 2006). Disruption of GARP proteins also causes defects in the transport of some glycosylphosphatidylinositol (GPI)-anchored and transmembrane proteins derived from the TGN, suggesting that GARP may have an important function in anterograde membrane trafficking (Hirata et al., 2015). In addition to endosomal trafficking defects, knockdown of the GARP subunits in mammalian cells can cause lysosome dysfunction (Perez-Victoria and Bonifacino, 2009; Perez-Victoria et al., 2008). Together, these data suggest that Vps54 plays an important role in endolysosomal trafficking pathways.

The glutamatergic *Drosophila melanogaster* larval neuromuscular junction (NMJ) is a well-characterized synapse that is used as a model system to study both neurodevelopmental and neurodegenerative processes (Deshpande and Rodal, 2016). In order to better understand the function of Vps54 in MNs, we have examined the development of the NMJ following depletion of the single *Drosophila* ortholog of *Vps54* (called “*scattered”* or “*scat”*). Here we show that both disruption of *scat* expression in an amorphic mutant and MN-specific reduction of expression via transgenic RNAi causes overgrowth of the larval NMJ. Unlike what is seen in yeast and mammalian cells, depletion of *scat* has no impact on the size, number, or localization of either early or late endosomes (EEs and LEs) (Palmisano et al., 2011; Quenneville et al., 2006). Both phenotypes are distinctly different from those reported in *Vps54* reduction-of-function (wobbler) and loss-of-function mouse models (Palmisano et al., 2011; Schmitt-John et al., 2005). The MN-specific knockdown of *scat* paired with overexpression of Rab5, Rab7, and Rab11, all suppress *scat* neurodevelopmental NMJ phenotypes. Conversely, presynaptic knockdown of *scat* combined with disruption of Rab7 function significantly decreases NMJ complexity and alters the composition of the postsynaptic density (PSD). This suggest that Vps54-mediated endosomal trafficking is required to control NMJ development and synaptic morphology in *Drosophila*.

## MATERIALS AND METHODS

### Drosophila genetics

The following lines were obtained from the Bloomington Stock Center: *P(PZ)scat^1^cn^1^, cn^1^, Df(2L)Exel8022, UAS-TRiP(HMS01910), UAS-LUC.VALIUM10, tub-Gal4, C380-Gal4, D42-Gal4, 24B-Gal4, UAS-YFP:Rab5, UAS-YFP:Rab5(S43N), UAS-YFP:Rab7, UAS-YFP:Rab7(T22N), UAS-YFP:Rab11, UAS-YFP:Rab11(S25N).* The *scat^1^cn^1^* and *cn^1^* lines were crossed into *w\** to normalize the genetic backgrounds*. w*; cn^1^* was used as a control to rule out any phenotypes that might be caused by *cn^1^* homozygosity in the *scat^1^* homozygote. The *Df(2L)Exel8022* line deletes the entire *scat* gene plus about 60 kb of flanking genomic DNA including 9 neighboring genes. The *Df* line does not contain the *cn^1^* allele. The *UAS-HA:scat* line was made by amplifying the *scat-RA* open reading frame from the LD22446 cDNA (Berkeley *Drosophila* Genome Project) with a 5’ primer containing the HA tag and then cloned into pUAST. The genomic rescue line (*scat-HA:scat*) was made by amplifying the *scat* cDNA and ∼350 nt of upstream and ∼50 nt of downstream genomic DNA with a 5’ primer containing the HA tag and cloned into pCASPR4. Both transgenic fly lines were generated by Bestgene. The *UAS-HA:scat* (*tub-Gal4>UAS-HA:scat*) and *scat-HA:scat* constructs rescued both *scat^1^* semi-lethality and male sterility (data not shown). All fly lines and crosses were maintained on standard Bloomington media in a diurnal 25°C incubator. Statistical analysis was done using larvae from each genotype raised under identical conditions.

### Immunohistochemistry and confocal microscopy

Larval body wall preps for NMJ and muscle analysis were dissected in Ca^2+^-free HL3 saline. Unless otherwise indicated, larvae were immunostained as previously described (Nesler et al., 2016). For imaging of the CNS, larval ventral ganglia and proximal axons were explanted and fixed in 4% paraformaldehyde in PBS. For GluRIIA and GluRIIB (and specific experiments with some Rab and DLG) antibodies, larvae were fixed with Bouin’s solution for 10 minutes. All were blocked in PBS containing 0.3% Triton X-100 (PBST), 2% BSA, and 5% normal goat serum for 30 minutes before incubation overnight with primary antibodies diluted in block. Following washes in PBST, CNS samples were incubated overnight with appropriate secondary antibodies.

All were mounted in DAPI Fluoromount G (Southern Biotech) for confocal microscopy. Primary antibodies used were anti-HA (1:1000) (Sigma; 3F10), Lva (1:50) (Sisson et al., 2000), Sxy16 (1:500) (Abcam; ab32340), Dlg (1:100) (DSHB; 4F3), Rab5 (1:800) and Rab11 (1:4000) (Tanaka and Nakamura, 2008), Rab7 (1:1500) (DSHB), GluRIIA (1:1000) and GluRIIB (1:1000) (Ramos et al., 2015), Brp (1:1000) (DSHB; nc82), and Dylight 649-conjugated anti-HRP (1:1000) (Jackson Labs). The Rab7 antibody was deposited to the DSHB by S. Munro (Riedel et al., 2016), Dlg by C. Goodman (Parnas et al., 2001), and Brp by E. Buchner (Wagh et al., 2006). Anti-mouse and rabbit secondary antibodies were conjugated to Alexa 488, 568, and 633 (Molecular Probes). All imaging was done on an Olympus FV1000 or FV3000 scanning confocal microscope with 40X, 60X, or 100X objectives (N.A. 1.30, 1.42, and 1.40 respectively). When shown, maximum Z projections were assembled from 0.4 μm optical sections using Olympus FV software. All image post-processing was done using Adobe Photoshop or ImageJ2 in open-source Fiji (Schindelin et al., 2012; Schindelin et al., 2015). For colocalization analysis, between 3 and 8 images were examined per experiment. Images were manually thresholded and the Pearson Correlation coefficients calculated using the JACoP plugin for ImageJ2/Fiji (Bolte and Cordelieres, 2006).

### Analysis of bouton number, synapse morphology, and active zones

The number of type 1 synaptic boutons was manually counted at muscles 6 and 7 (m6/7) in abdominal segment 3 (A3) as previously described (Pradhan et al., 2012). To account for differences between genotypes in the scaling of NMJs to muscle size, synaptic bouton numbers were normalized to muscle surface area (MSA). MSA was calculated using ImageJ2/Fiji from images of m6/7 obtained using a 20X objective (N.A. 0.85). Branching was determined by counting branch points between strings of boutons at least 3 boutons long. The same NMJs were subjected to analysis using the Morphometrics algorithm, A Fiji-based macro that quantifies morphological features of *Drosophila* synapses (Nijhof et al., 2016). The parameters examined here include total bouton counts, NMJ area, and NMJ length. To validate these results, we compared bouton number determined by the macro with manual counts for *scat* mutant analysis. While total bouton numbers were not identical, macro counts correlated significantly with manual counts (Pearson correlation coefficient = 0.76; C.I. 95% 0.66-0.84; p < 0.0001; n = 91 NMJs). Analysis was done using Fiji version 2.0.0 and the NMJ Morphometrics plugin version 20161129. Settings used were: maxima noise tolerance = 500, small particle size = 10, minimum bouton size = 10, and rolling ball radius = 500. NMJ outline and skeleton thresholds were set to “triangle”.

To quantify active zone number, NMJs were counterstained with antibodies targeting Brp and HRP as described above. Maximum Z-projections were processed using the TrackMate plugin for ImageJ2/Fiji (Tinevez et al., 2017). Active zone images were opened in TrackMate using the default calibration settings for the Brp channel. The following additional settings were used: LoG detector = on, estimated blob diameter = 1 μm, and threshold settings = 100. The results were previewed to ensure accurate detection of found spots and data recorded for all boutons.

### Behavioral analysis

For the analysis of larval crawling, videos were collected using an iPhone XR (Apple) set in time-lapse video mode (2 frames per second). Ten larvae were collected for each genotype and transferred to the center of a room temperature 15 cm petri dish containing 2% agarose. 45-90 second videos of each larva were collected in triplicate. Videos were trimmed to 90 frames using the Apple photo editing trimming tool selecting for direct path larval movement away from the edge of the petri dish. Files were then converted from .MOV to .TIF series using the export function in ImageJ2/Fiji. Subsequent analysis was done using TrackMate, a Fiji-based macro developed for single particle tracking (Tinevez et al., 2017). Images were adjusted to maximize contrast between larva and the background. The parameters were adjusted as follows to analyze larval locomotion. Settings used were: LoG detector, HyperStack displayer, simple LAP tracker, and spot tracking were all turned on. The average velocity was recorded for each replicate.

### Statistics

All data was recorded in Excel (Microsoft) and graphed and analyzed in Prism (GraphPad). Results were considered to be statistically significant at p < 0.05. Results shown throughout the study are mean ± SEM. n.s. = not significant, * p < 0.05, ** p < 0.01, *** p < 0.001, and **** p < 0.0001. Data for *scat^1^* loss of function and larval crawling velocity were both analyzed by Kruskal-Wallis followed by a Dunn’s multiple comparison test to determine significance. Each *scat* RNAi experiment had its own control and was analyzed using a Mann-Whitney U test. The number of synaptic boutons and Brp-positive AZs in genetic interaction experiments where both analyzed by one-way ANOVA followed by a Holm-Sidak multiple comparison test.

## RESULTS

### *scat* is required to control axon terminal growth at the larval NMJ

To determine whether *scat* has a function in fly MNs, we examined the development of a well-characterized NMJ in third instar larvae. The classic *scat^1^* allele is a P-element insertion near the 5’ end of the second coding exon of the *scat* gene (Figure S1) (Castrillon et al., 1993). Protein expression is completely disrupted in *scat^1^* homozygotes suggesting that it is a null allele (Fari et al., 2016b). We found that the morphology of the NMJ was distinctly different in *scat^1^* mutants compared to controls (Figure 1A). Quantitative analysis of the number of type 1 synaptic boutons revealed a greater than 2-fold overelaboration in *scat^1^* null animals (Figure 1B; 114% increase; p < 0.0001). A similar phenotype was observed when the *scat^1^* allele was placed in *trans* to the overlapping *Df(2L)Exel8022* deficiency (Figures 1A-B; 94% increase; p < 0.0001). Quantification of the number of synaptic arbor branch points correlated strongly with synaptic boutons (Figure 1C). The *scat^1^* homozygous mutant phenotype was rescued when a transgenic construct was introduced back into the *scat^1^* background (Figures 1A-C). This rescue transgene includes the *scat* promoter and ∼350 bp of upstream genomic DNA regulating the expression of a hemagglutinin (HA)-tagged *scat* cDNA (*scat-HA:scat*). Taken together, these data suggest that *scat* has a critical function in the control of axon terminal growth during larval NMJ development.

During our NMJ analyses, we noted that synaptic bouton morphology appeared to also be abnormal in *scat^1^* mutants. The NMJ at muscle 6/7 has two types of synaptic boutons – type 1b (big) and type 1s (small) boutons, which are derived from two distinct MNs and differ in morphology and physiology (Menon et al., 2013). Immunostaining of Discs large (Dlg), the fly ortholog of the postsynaptic density protein, PSD-95, is also usually stronger in in type 1b boutons and weaker in type 1s boutons (Lahey et al., 1994). In contrast to controls, Dlg staining in *scat^1^* homozygotes is roughly the same in both types of boutons and is spotty and discontinuous suggesting that localization of Dlg to postsynaptic densities may be partially disrupted (Figure 1D). Additionally, both types of boutons were smaller in *scat^1^* mutants (Figure 1D). Quantification of this revealed a significant 2-fold decrease in synaptic area per bouton and an increase in total synaptic length in *scat^1^* homozygotes (Figure 1E-F; area = 50% of controls; p < 0.0001; length = 173% of controls; p < 0.0001). In contrast, a robust a reduction in bouton size was not seen in *scat^1^/Df* larvae (Figure 1E). However, the deficiency line deletes the *scat* gene region plus about 60 kb of flanking sequence including nine neighboring genes. It is possible that heterozygosity of one or more of these loci might have an uncharacterized effect on NMJ growth. Both the “small bouton” and length phenotypes were rescued by the *scat-HA:scat* transgene (Figures 1E-F).

**Figure 1.**
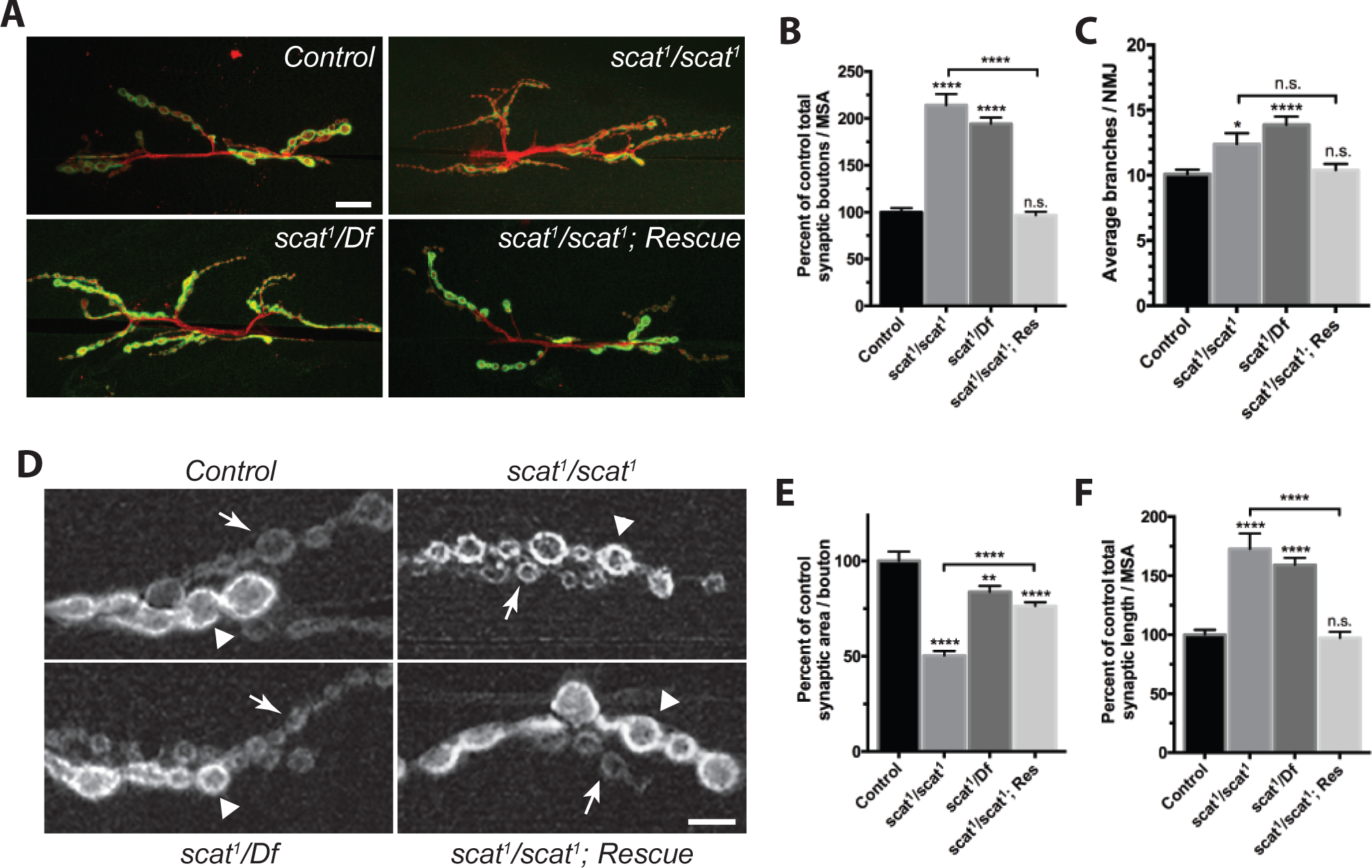
*scat* is a negative regulator of synaptic development at the larval NMJ. (A) *scat* loss-of-function causes defects in NMJ structure. Wandering third instar larvae from controls, *scat^1^* homozygotes, *scat^1^/Df(2L)Exel8022*, and the *scat^1^/scat^1^; scat-HA:scat/scat-HA:scat* rescue (Res) lines were stained with antibodies targeting the postsynaptic density marker, Dlg (green) and the neuronal membrane marker, HRP (red). Images show maximum Z-projections. The NMJs innervating body wall muscles 6/7 in abdominal segment 3 (m6/7 in A3) were analyzed. *scat* mutants have an increased number of boutons and synaptic arbors in comparison to controls. Scale bar, 20 μm. (B) Total bouton number/MSA (normalized to control) and (C) synapse branch points are significantly increased in *scat* mutants. Both were quantified by counting manually and both phenotypes are rescued by the introduction of the *scat-HA:scat* transgenic construct. *N* = 23, 21, 24, and 25. (D) *scat* loss-of-function causes defects in the size of both type 1b (arrowhead) and 1s boutons (arrows). Wandering third instar larvae from controls, *scat^1^* homozygotes, *scat^1^/Df(2L)Exel8022*, and the *scat^1^/scat^1^; scat-HA:scat/scat-HA:scat* rescue lines were stained with an antibody targeting Dlg. Images shown are single focal planes through the equator of the type 1b boutons. *scat^1^* homozygotes have noticeably smaller boutons than controls. The NMJs innervating muscle 6/7 in body segment A3 were analyzed. Scale bar, 5 μm. (E) Total synaptic length/MSA (normalized to control) is significantly increased and (F) synaptic area per bouton is decreased in *scat^1^* homozygotes. Both features were quantified using the Morphometrics algorithm. *N* = 23, 19, 24, and 25. Data are represented as the mean ± SEM. Unless otherwise indicated, all comparisons have been made to the control. * p < 0.05, ** p < 0.01, **** p < 0.0001.

### *scat* function is required in both the MN and muscle to control NMJ development

Given the critical role of *Vps54* in mouse MNs, we postulated that *scat* might have an important presynaptic function at the larval NMJ. To investigate this, we targeted the knockdown of *scat* expression in MNs using a transgenic short hairpin RNA (shRNA) driven by the MN-specific *C380-Gal4* driver. Presynaptic knockdown of *scat* resulted in a highly significant increase in the number of boutons (Figures 2A-B; 81% increase; p < 0.0001). To confirm these results, we drove expression of the shRNA transgene using a second, albeit weaker, MN driver, *D42-Gal4*, and observed a similar increase in the number of synaptic boutons (Figure 2B; 37% increase; p < 0.0001). To determine if the function of *scat* was restricted to the MN, we examined targeted knockdown of *scat* expression in larval muscle using the muscle-specific driver *24B-Gal4* (Figures 2A-B). Knockdown in the postsynaptic muscle also resulted in a small but significant increase in bouton number relative to controls (Figure 2B; 19% increase; p = 0.0207). As in *scat^1^* mutants presynaptic knockdown of *scat* also caused a significant increase in the number of synaptic arbor branches (Figure 2C; *C380>scat shRNA*; p < 0.0001). Collectively, these data suggest that *scat* functions in both the pre- and post-synaptic compartments to control NMJ development.

**Figure 2.**
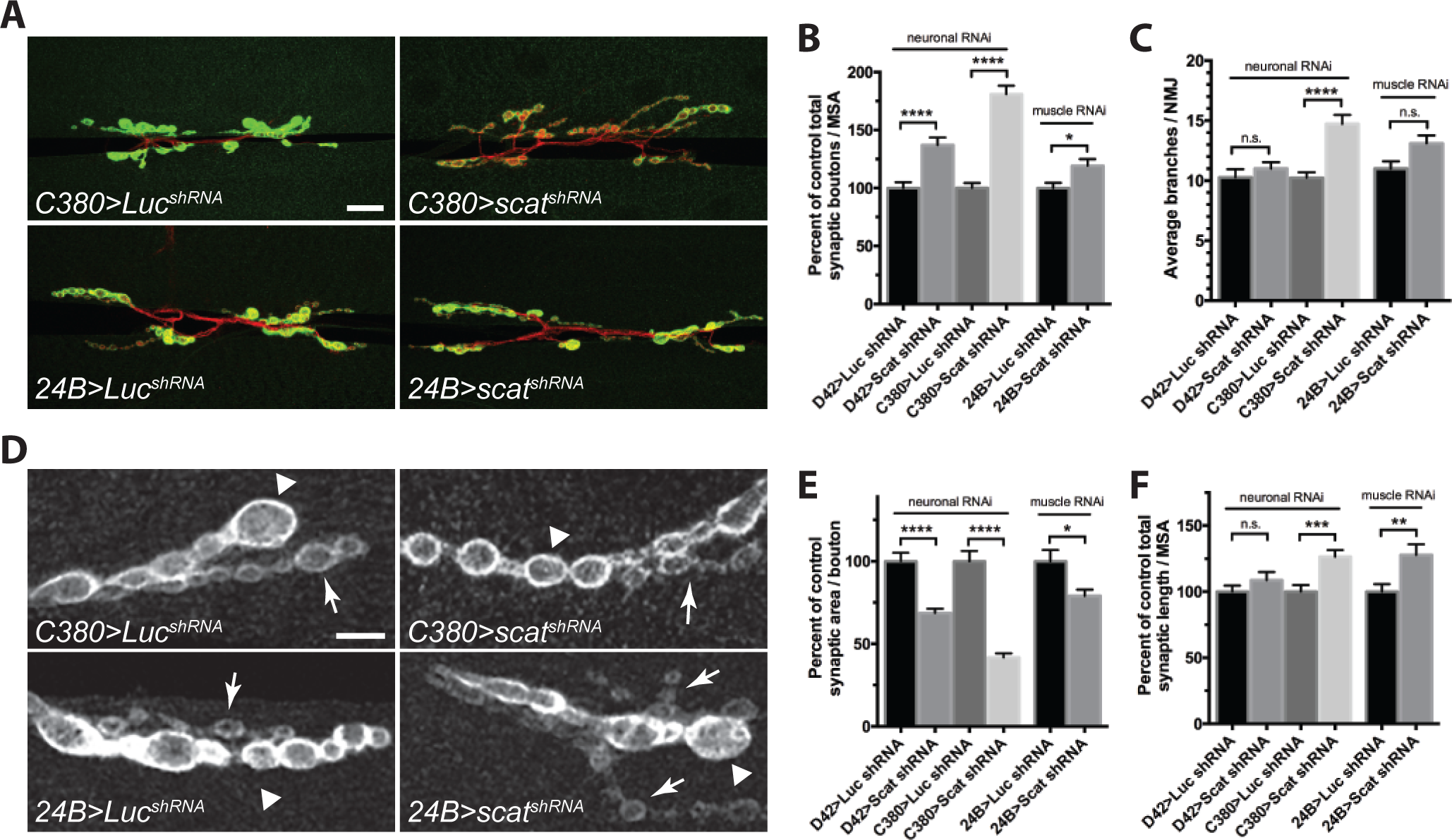
*scat* has a strong presynaptic function in the control of NMJ development. (A) Knockdown of *scat* expression in the presynaptic motor neuron by RNAi causes defects in NMJ structure. An inducible transgenic shRNA targeting luciferase (*UAS-LUC.VALIUM10*) or *scat* (*UAS-TRiP^HMS01910^*) was expressed in motor neuron using the *C380-Gal4* or the weaker *D42-Gal4* driver (*D42* not shown) or in muscle using *24B-Gal4*. NMJs at muscle 6/7 in body segment A3 in late third instar larvae were stained with antibodies targeting Dlg (green) and HRP (red). Images show maximum Z-projections. Presynaptic knockdown of *scat* causes an increased number of boutons and synaptic arbors. Scale bar, 20 μm. (B) As in *scat* mutants, the total bouton number/MSA (normalized to the respective control) and (C) synapse branch points are significantly increased by presynaptic *scat* knockdown. *N* = 18, 18, 17, 18, 20 and 25. (D) Presynaptic *scat* knockdown causes defects in the size of type 1b (arrowhead) and 1s boutons (arrows). Wandering third instar larvae from genotypes indicated in A were stained with an antibody targeting Dlg. Images show single focal planes through the equator of type 1b boutons. Presynaptic *scat* knockdown causes a reduction in the size of type 1 boutons. Scale bar, 5 μm. E) As in *scat* mutants, total synaptic area per bouton/MSA (normalized to control) is significantly decreased and (F) length is increased by presynaptic *scat* knockdown. Effects are statistically significant but not as dramatic following *scat* RNAi in the postsynaptic muscle. Both features were quantified using the Morphometrics algorithm. *N* = 18, 17, 17, 18, 20, and 19. All statistical comparisons shown have been compared to driver-specific controls (*driver/+* heterozygotes). Data represented as the mean ± SEM. * p < 0.05, ** p < 0.01, *** p < 0.001, **** p < 0.0001.

We were also interested to see if bouton morphology was altered by pre-or postsynaptic disruption of *scat* expression. While the small bouton phenotype was not as obvious in either case (Figure 2D), presynaptic RNAi does significantly reduce bouton size. *C380>scat shRNA* caused a 58% reduction in synaptic area per bouton (Figure 2E; p < 0.0001). A similar result was observed using the *D42-Gal4* driver (Figure 2E; 41% decrease; p < 0.0001). *C380>scat shRNA* also had a modest but significant effect on total NMJ length (Figure 2F; 27% increase; p = 0.0006). As seen in *scat^1^* mutants, knockdown of *scat* in motor neurons (but not in muscle) partially disrupted normal Dlg expression and localization to post-synaptic densities (Figure 2D). Disruption of *scat* expression in the muscle also had a significant effect on average bouton size (Figure 2E; 21% decrease; p = 0.0303) and length of NMJs (Figure 2F; 28% increase; p = 0.0043). Together, these data suggest that *scat* is required on both sides of the synapse to control bouton morphology.

Conversely, we asked if *scat* overexpression impacted NMJ development. To test this, we constructed a fly line containing a Gal4-inducible version of the HA-tagged *scat* cDNA (*UAS-HA:scat*). The global overexpression of HA:Scat in a wild-type background (*tubulin-Gal4>UAS-HA:scat*) had no impact on NMJ development (Figure S2). Neuronal expression of *UAS-HA:scat* with a strong pan-neuronal driver (*elav-Gal4*) caused a small but statistically significant increase in branching and synaptic area per bouton (Figures S2D-E; branching, p = 0.0159; synaptic area, 21% increase; p = 0.0008). In contrast, overexpression using a strong muscle driver (*Mef2-Gal4*) resulted in a significant increase in branch and synaptic bouton number with no changes in bouton morphology (Figures S2A-D; branching, p = 0.0072; bouton number, 25% increase, p = 0.0003). Together, these data suggest that both loss- and gain-of-function of *scat* causes NMJ defects.

### Scat localizes to the *trans-*Golgi in motor neurons, glia, and muscle cells

Vps54 localizes primarily to the TGN in yeast and in mouse spermatids (Berruti et al., 2010; Conibear and Stevens, 2000). Similar localization patterns have been observed with fluorescently-tagged Scat protein in *Drosophila* (Fari et al., 2016b). Based on this, we predicted that Scat would localize to the TGN in larval MNs and muscle cells. In our hands, the only available antibody targeting Scat did not work in neurons for immunofluorescence. Moreover, HA-tagged Scat expression from two copies of the *scat-HA:scat* transgene could not be detected. Therefore, to examine subcellular localization, we drove expression of *UAS-HA:scat* using the ubiquitous *tub-Gal4* driver and labeled the HA:Scat protein using an antibody targeting the HA tag. First, we counterstained with an antibody that recognized the golgin Lava lamp (Lva), a marker for the *cis-* Golgi (Sisson et al., 2000). We found that most HA:Scat was juxtaposed to Lva staining in the soma of larval MNs (Figure 3A), the ensheathing glial cells surrounding peripheral nerves (Figure 3B), and in body wall muscle (Figure 3C). Both HA:Scat and Lva was absent from motor axons (blue in Figure 3B) and presynaptic boutons (grey in Figure 3C). Taken together, these data are consistent with HA:Scat localizing to a structure in close proximity to the *cis-*Golgi.

**Figure 3.**
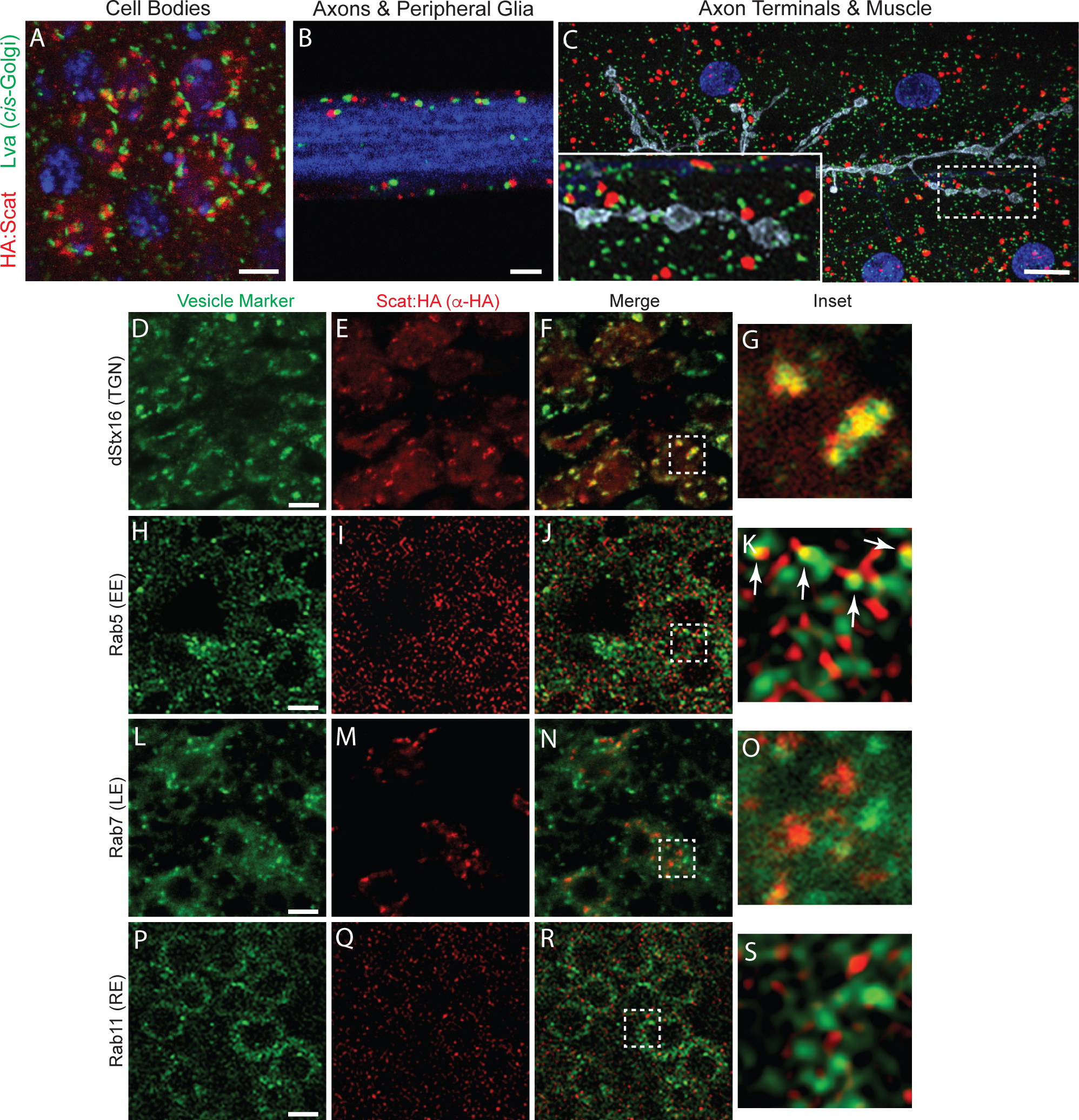
Scat localizes to perikaryon but not peripheral axons or axon terminals. (A-C) Scat localizes to a structure adjacent to the *cis*-Golgi in (A) motor neuron cell bodies, (B) peripheral glia, and (C) body wall muscle. (A-B) Ventral ganglia and (C) body wall muscle preps from wandering third instar larvae expressing inducible *HA:scat* under control of the *tubulin-Gal4* driver were stained with antibodies targeting the HA tag (red) and Lva (green). Single focal planes are shown in A while B and C are maximum Z-projections. HA:Scat localizes to the motor neuron cell body but not peripheral axons or axon terminals. Most HA-positive structures are adjacent to the Lva-positive *cis-*Golgi. Blue is DAPI (DNA) in A and Hrp (axon) in B. Grey in C is Hrp (axon). Scale bars are 2.5 μm in A and 10 μm in B and C. (D-S) *tub-Gal4>HA:scat* animals were counterstained with antibodies targeting the HA tag (red) and the indicated marker (green). Images shown are single focal planes. Vesicle trafficking markers shown are the TGN marker, Syntaxin 16 (D-G), the early endosome marker, Rab5 (H-K), the late endosome marker, Rab7 (L-O), and the recycling endosome marker (P-S). Boxed areas indicated in the merged images (F, J, N, and R) are shown in G, K, O, and S (respectively). Scale bars in D, H, L, and P are 2.5 μm.

Because of the strong presynaptic role for *scat* in the control of NMJ development, we focused on further characterizing the localization of Scat within MNs. To confirm localization of Scat to the TGN, we counterstained with an antibody targeting *Drosophila* Syntaxin-16 (dStx16), a core component of the t-SNARE that is involved in retrograde transport of vesicles derived from EEs and LEs to the TGN (Amessou et al., 2007; Quenneville et al., 2006). As expected, HA:Scat colocalized strongly with dStx16-positive foci in the MN cell bodies (Figures 3D-G; Pearson’s correlation coefficient (PCC) = 0.55 ± 0.02). This is consistent with the localization of Scat to the TGN and its role in recruiting and assembling t-SNAREs at this membrane (Perez-Victoria and Bonifacino, 2009). In yeast, Vps54 localizes to EEs via a conserved C-terminal domain and it is required for retrograde transport from EEs to the TGN (Quenneville et al., 2006). To examine the colocalization of HA:Scat with endosomal compartments, we counterstained larval MNs with antibodies targeting the small GTPases, Rab5, Rab7, and Rab11, which are used as markers for EEs, LEs, and recycling endosomes (REs) respectively. In contrast to dStx16, we observed little colocalization between HA:Scat and Rab5, Rab7, or Rab11 (Figures 3H-S; PCC = 0.15 ± 0.01, 0.22 ± 0.04, and 0.178 ± 0.02). Interestingly, HA:Scat sometimes appeared to localize to a structure immediately adjacent to Rab5-positive EEs (arrows in Figure 3K). A similar result was not observed with either Rab7- or Rab11-postive endosomes (Figures 3O and 3S).

### Localization of Syntaxin-16 to the TGN is disrupted in *scat* mutant motor neurons

Endosomal trafficking is required for the bi-directional transfer of membranes and receptors along axons and dendrites, which is likely to be important in regulating different aspects of synaptic development (Lasiecka and Winckler, 2011). To begin to understand how loss of *scat* expression in larval MNs leads to synaptic overgrowth, we analyzed the impact on the localization of markers for endocytic trafficking pathways. In yeast and mammalian cells, GARP is required to tether vesicles derived from both EEs and LEs to the TGN (Conboy and Cyert, 2000; Conibear and Stevens, 2000). It does so, in part, by controlling the assembly of the t-SNARE complex (Perez-Victoria and Bonifacino, 2009). Based on this, we asked if dStx16 localization would be disrupted in *scat* loss- or reduction-of-function mutant MNs. As predicted, dStx16 antibody staining was diffuse in *scat^1^* homozygous larvae (Figure 4A), which was rescued by the introduction of the *scat-HA:scat* transgene. Very similar results were seen following targeted disruption of *scat* expression in larval MNs by RNAi (Figure S3A). Some punctate dStx16 fluorescence remains in *scat* mutants and after Scat RNAi suggesting that this phenotype is only partially penetrant. Together, these data indicate that *scat* contributes to dStx16 localization or membrane association at the TGN. Moreover, this suggests that an integral component of the retrograde trafficking pathway (the t-SNARE complex) has been at least partially disrupted.

**Figure 4.**
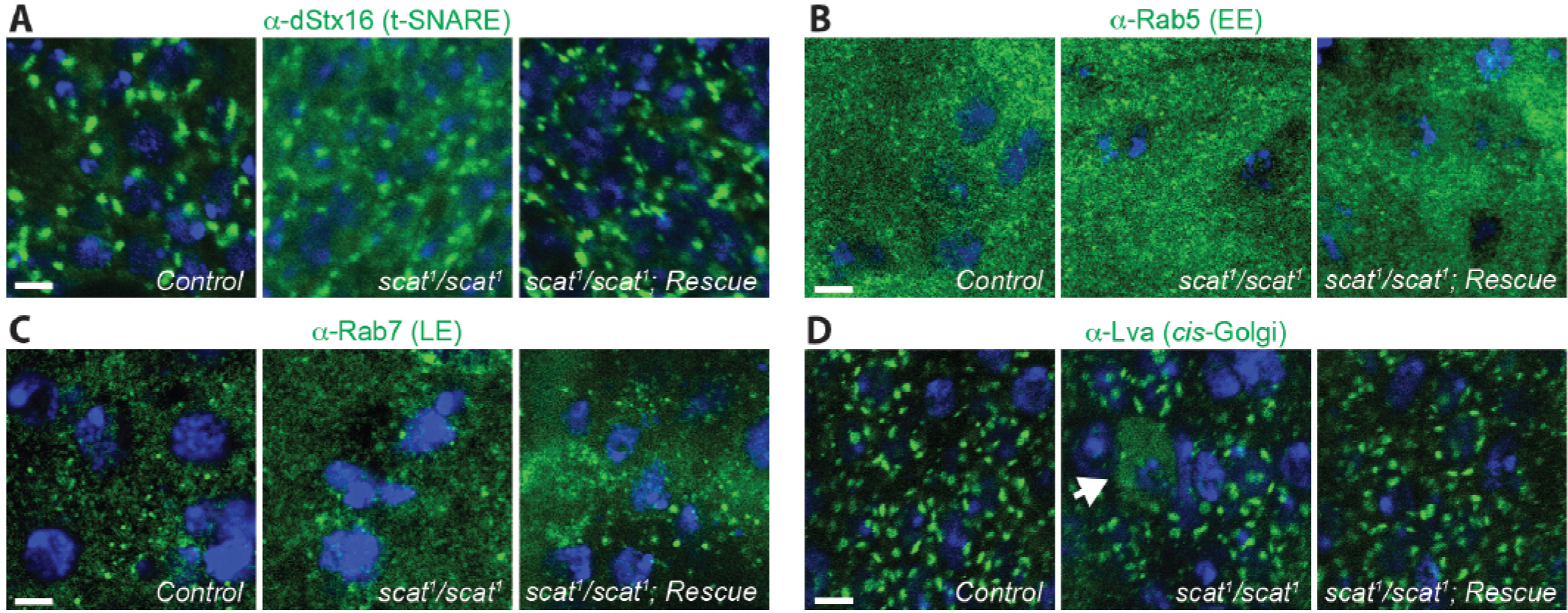
*scat* mutant motor neurons have defects in Syntaxin-16 localization and *cis*-Golgi integrity. (A) Localization of Syntaxin-16 to the TGN is disrupted in *scat^1^* mutants. Images shown are single focal planes. Ventral ganglia from wandering third instar larvae controls, *scat^1^* homozygotes, and the *scat^1^/scat^1^; scat-HA:scat/scat-HA:scat* rescue lines were stained with an antibody targeting dStx16 (green) and DAPI to visualize nuclei (blue). dStx16 staining is significantly more diffuse (but still clearly punctate) in *scat^1^* mutants compared to controls. This phenotype is rescued by the introduction of the *scat-HA:scat* transgene. (B) EEs are not affected in *scat^1^* mutants. Indicated genotypes have been stained with an antibody targeting Rab5 (green) and DAPI (blue). (C) LEs are not affected in *scat* mutants. Indicated genotypes have been stained with an antibody targeting Rab7 (green) and DAPI (blue). (D) Localization of the *cis-*Golgi marker, Lva is partially disrupted in the cell body of some *scat* mutant motor neurons (arrows). The indicated genotypes have been stained with an antibody targeting Lva (green) and DAPI (blue). This phenotype is never observed in control or rescue larvae. Scale bar, 2.5 μm.

Next, we determined if *scat* loss-of-function had an impact on endosomal pools in MNs. Disruption of *Vps54* expression in yeast causes the accumulation of vesicles containing markers for EEs (Quenneville et al., 2006). In contrast, MNs in the wobbler mouse appear to accumulate large Rab7-positive LEs (Palmisano et al., 2011). To determine if endosomal trafficking is affected by the disruption of *scat* expression, we examined the localization of markers for EEs and LEs in larval MNs. Surprisingly, neither the size or number of Rab5- and Rab7-postive endosomes were obviously altered in *scat^1^* homozygous mutant larvae (Figure 4B-C). Thus, at least at this stage of development, disruption of *scat* does not have a significant impact on endosomal populations.

Although *Vps54* loss-of-function in cultured mammalian cells causes defects in vesicle trafficking pathways, there is no apparent impact on Golgi structure or function (Karlsson et al., 2013). In contrast, MNs in the wobbler mouse show signs of Golgi dysfunction and fragmentation beginning in early stages of neurodegeneration (Palmisano et al., 2011). In order to examine Golgi structure after the disruption of *scat* expression, we stained larval MNs with antibodies targeting Lva (Sisson et al., 2000). Interestingly, we found that some MN cell bodies in *scat^1^* homozygotes and *scat^1^/Df(2L)Exel8022* larvae had diffuse and cytoplasmic Lva staining suggesting that *cis-*Golgi integrity in some neurons has been at least partially disrupted (Figure 4D). We never observed this phenotype in controls or transgenic rescue animals (Figure 4D). We observed a similar phenotype following the targeted depletion of *scat* in larval MNs by RNAi (Figure S3B). However, Lva staining was more globally diffuse and punctate structures remained much more intact in affected neurons. Collectively, these data suggest that *scat* expression is required, at least in part, to maintain Golgi integrity in the perikaryon of larval MNs.

### *scat* interacts genetically with Rab proteins to control axon terminal growth

We were next interested in gaining some mechanistic understanding into how the disruption of *scat* expression in MNs leads to the overgrowth of axon terminals. Rab proteins not only associate with specific endosomal compartments, their activity is also required to mediate all steps of membrane trafficking (Pfeffer, 2017). Rabs function by switching between GDP- and GTP-bound forms which regulates their ability to bind to specific Rab effector proteins (Pfeffer and Aivazian, 2004). Transgenic *Drosophila* lines have been constructed that contain Gal4-inducible Rabs that are GTP-binding defective conferring dominant-negative activity and allow for the cell autonomous disruption of Rab function (Zhang et al., 2007). In our hands, MN-specific expression of wild-type and dominant-negative Rab5, Rab7, and Rab11 had no effect on the number of type 1 synaptic boutons at larval muscle 6/7 compared to controls (Figure 5A-B; data not shown). If *scat* interacts genetically with Rab proteins to control synaptic development, we expected that co-expression of wild-type or dominant-negative Rabs would enhance or suppress the *scat* knockdown phenotype. Interestingly, co-expression of both forms of Rab5, Rab7, and Rab11 significantly suppressed the *scat* shRNA overgrowth phenotype. In most cases, these NMJs appeared to be morphologically indistinguishable from negative controls (Figure 5A; data not shown). There were two notable exceptions to this. First, co-expression of wild-type Rab5 with the *scat* shRNA often caused the formation of clusters of synaptic boutons instead of the normal “beads on a string” phenotype (Figure 5C). Second, in *C380>scat shRNA, Rab7 (DN)* animals, postsynaptic Dlg staining appears to be appreciably disrupted (Figure 5C). Moreover, total bouton number was significantly reduced when compared to *C380>scat shRNA* and *C380>Rab7 (DN)* controls (54% decrease, p < 0.0001 and 46% decrease, p < 0.0001 respectively). Taken together, these data suggest that the regulation of normal NMJ development by Scat requires endosomal trafficking mediated by Rab5, Rab7, and Rab11 activity. Disruption of both *scat* and *rab7* function in motor neurons significantly reduced NMJ complexity and triggers synapse disassembly.

**Figure 5.**
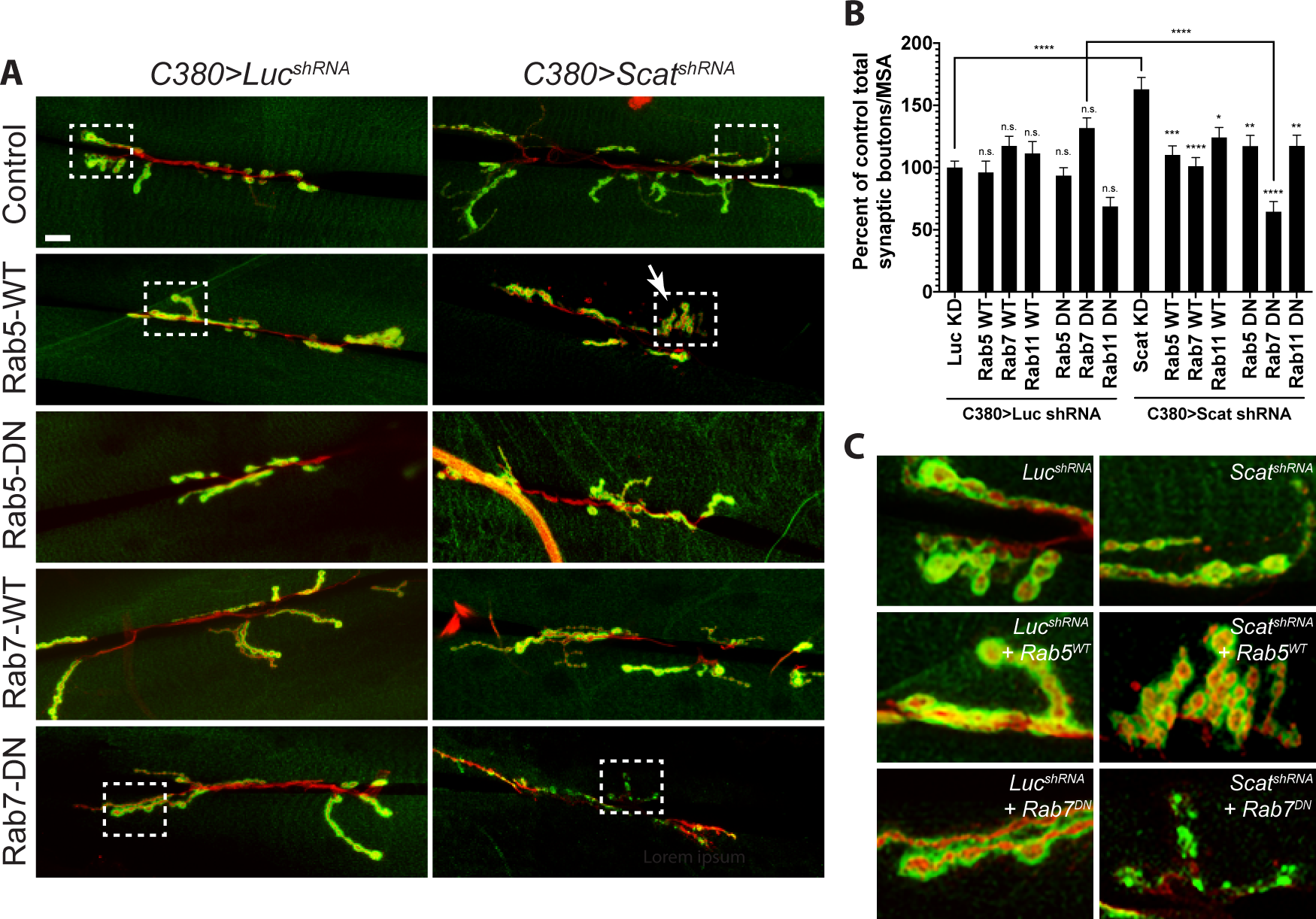
NMJ phenotypes caused by presynaptic *scat* knockdown are suppressed by Rab5, Rab7, and Rab11. (A) NMJ phenotypes caused by the motor neuron-specific knockdown of *scat* expression by RNAi are suppressed by overexpression of wild-type and dominant-negative transgenes for Rab5, Rab7, and Rab11 (Rab11 images are not shown). An inducible transgenic shRNA targeting luciferase (*UAS-LUC.VALIUM10*) or *scat* (*UAS-TRiP^HMS01910^*) was expressed in motor neuron using the *C380-Gal4* driver in combination with an inducible YFP-tagged wild-type or dominant negative Rab5, Rab7, or Rab11 (*UAS-YFP:Rab*). NMJs at muscle 6/7 in body segment A3 in late third instar larvae were stained with antibodies targeting Dlg (green) and HRP (red). Images show maximum Z-projections. The boxed areas are blown up in C to show altered bouton or PSD morphologies. Scale bar, 20 μm. (B) As shown in Figure 2, the total bouton number/MSA (normalized to the respective control) are significantly increased by presynaptic *scat* knockdown. This phenotype is suppressed by co-expression of wild-type and dominant negative Rabs. *C380>scat shRNA, Rab7 (DN)* double mutant NMJs are significantly smaller. *N* = 18, 12, 16, 13, 22, 17, 21, 17, 18, 20, 20, 22, 21, and 23. (C) Boxed areas in A. Many *C380>scat shRNA, Rab5 (WT)* NMJs have a clustered bouton phenotype similar to many endocytic mutants. Dlg staining and bouton morphology is significantly disrupted in *C380>scat shRNA, Rab7 (DN)* double mutants. Data are represented as the mean ± SEM. Unless otherwise indicated, all comparisons have been made to the control. * p < 0.05, ** p < 0.01, *** p < 0.001. **** p < 0.0001.

### *scat* interacts genetically with *rab7* to regulate synapse integrity at the larval NMJ

Postsynaptic Dlg staining appears to be partially disrupted in *scat^1^* mutants and following presynaptic knockdown by RNAi (Figures 1D and 2D) and disruption of both *scat* and *rab7* in MNs significantly enhances this phenotype (Figure 5C). At the *Drosophila* NMJ, Dlg forms a multimeric scaffold that is required for the clustering of postsynaptic glutamate receptors (GluRs) containing the GluRIIB subunit but mutations in *dlg* have no effect on the localization of its alternative subunit, GluRIIA (Chen and Featherstone, 2005). Thus, we asked if GluR localization to postsynaptic sites was altered following the disruption of both *scat* and *rab7* function. High resolution, single focal plane images of NMJs confirmed that Dlg staining was reduced and spotty in type 1b synapses in *C380>scat shRNA, Rab7 (DN)* larvae compared to controls (Figure 6A). Strikingly, staining with GluRIIB antibodies was slightly reduced in *C380>scat shRNA* animals and significantly disrupted in *C380>scat shRNA, Rab7 (DN)* larvae (Figure 6B). In contrast, the localization of the GluRIIA subunit does not appear to be affected (Figure 6C) suggesting that the core GluR has not been lost from postsynaptic sites. This was confirmed by analyzing the localization of the requisite GluR subunit, GluRIIC which was similarly not affected (data not shown). Collectively, these data suggest that the Scat protein is required in the presynaptic cell to regulate the localization of Dlg and GluRIIB to postsynaptic sites via a mechanism that involves the activity of Rab7.

**Figure 6.**
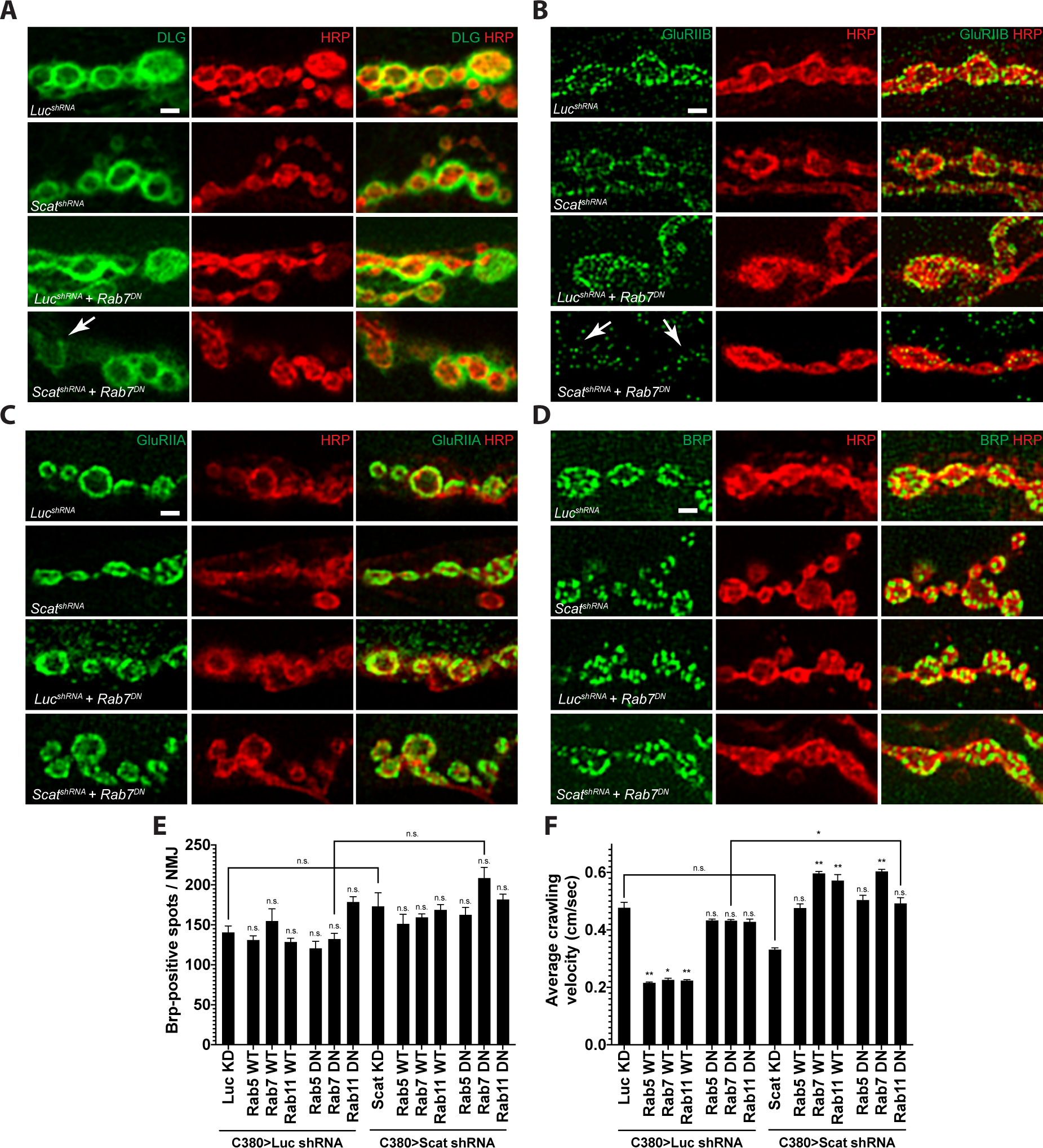
*scat* interacts genetically with *rab7* to control the composition of the PSD. Localization of the PSD proteins (A) Dlg, (B) GluRIIB, (C) GluRIIA, and AZ marker (D) Brp to synaptic boutons at muscle 6/7 in body segment A3 in late third instar larvae are shown in green. All NMJs have been counterstained with an antibody targeting HRP (red). Merged images are included to confirm pre- or post-synaptic localization. All images are single focal planes. Synaptic bouton structure has been better preserved here using Bouin’s fixative. The arrow in (A) points to a type 1b bouton with abnormally low levels of Dlg – neighboring boutons have spotty Dlg staining. Arrows in (B) point to synaptic boutons where GluRIIB localization has been significantly disrupted (compare *scat^shRNA^, Rab7^DN^* to any of the control genotypes). Scale bar, 2.5 μm. (E) Quantification of the number of Brp-positive spots per NMJ. No significant difference was observed in any genotype (*N* = 5 each). (F) Average crawling velocity of third instar larvae for each genotype (*N =* 10 each). MN-specific overexpression of wild-type Rab5, Rab7, and Rab11 alone significantly suppressed larval crawling velocity while overexpression of dominant negative Rabs had no effect. Interestingly, Overexpression of both wild-type and dominant negative forms of all Rabs significantly suppressed the *C380>Scat shRNA* phenotype. * p < 0.05, ** p < 0.01.

In order to further explore the role of *scat* in synaptic architecture at the NMJ, we next asked if *scat* was required to control the localization of Bruchpilot (Brp) to presynaptic boutons. Brp is similar to the human ELKS/CAST family of active zone (AZ) proteins and is required to regulate AZ structure and function at synapses (Wagh et al., 2006). At the larval NMJ, Brp localizes to presynaptic AZs and the lack of Brp causes defects in AZ assembly, Ca^2+^ channel clustering, and vesicle release (Kittel et al., 2006). We observed no significant difference in the total number of Brp puncta per NMJ in *C380>scat shRNA, Rab7 (DN)* larvae or any other genotype tested (Figure 6D-E). Similar results were observed in *scat^1^* homozygotes compared to controls and transgenic rescues (Figure S4). Instead, we observed a significant decrease in the number of AZs per synaptic bouton, which was partially rescued by the *scat-HA:scat* transgene (Figure S4; 7.0 ± 0.70 in controls vs. 4.5 ± 0.46 in *scat^1^* homozygotes, p = 0.009; *scat^1^* homozygotes vs. 5.1 ± 0.41 in *scat^1^* homozygotes with two copies of the *scat-HA:scat* transgene, p = 0.037). These results suggest that presynaptic AZs have not been disrupted.

Because disruption of *scat* expression (either alone or in combination with expression of a dominant-negative *rab7*) causes abnormal NMJ development, we next asked if larval crawling behavior was affected. Defects in larval locomotion have been directly linked to neuronal and synaptic dysfunction (Folwell et al., 2010; Mudher et al., 2004). Comparison of larval crawling speeds showed no statistical difference between *C380>scat shRNA* animals and controls (Figure 6F). Interestingly, MN-specific expression of wild-type Rab5, Rab7, and Rab11 all significantly decreased larval crawling velocity (55% decease, p = 0.0044; 53% decrease, p = 0.0173; and 53% decrease, p = 0.0184 respectively). However, expression of the dominant-negative forms of these proteins alone had no effect (Figure 6F). Conversely, MN-specific knockdown of *scat* along with expression of wild-type Rab7, Rab11 and dominant negative Rab7 all significantly increased average crawling velocity compared to *C380>scat shRNA* controls (44% increase, p < 0.0001; 72% increase, p = 0.0013; and 48% increase p < 0.0001, respectively). Thus, there appears to be no correlative relationship between larval crawling behavior and either synapse morphology or post-synaptic density composition in *C380>scat shRNA* and *rab* transgenic animals.

## DISCUSSION

The GARP complex is required to control a wide range of cellular processes highlighting the importance of understanding more about its function (Bonifacino and Hierro, 2011). Notably, the disruption of two subunits of GARP have been linked to neuronal dysfunction in mammals: 1) a homozygous recessive mutation in *Vps54* in the wobbler mouse (Schmitt-John et al., 2005), and 2) heterozygous mutations in *Vps53* in pontocerebellar hypoplasia type 2E in humans (Feinstein et al., 2014). Vps54 associates specifically with the GARP complex while Vps53 is a core component of both GARP and the endosome-associated protein (EARP) complex (Schindler et al., 2015). An increased number of swollen Rab7-positive LEs have been reported in both cases. Here, we show that the loss of *scat* expression in *Drosophila* MNs appears to have no impact on LE size or number (Figure 4C). However, presynaptic *scat* interacts genetically with *rab7* to control NMJ growth and composition of the PSD (Figures 5 and 6). These data strongly suggest that GARP-mediated LE to TGN retrograde transport is critical for NMJ development.

Our results demonstrate that *Drosophila* Vps54 localizes to the TGN (Figures 3D-G) and that the localization of dStx16, a core TGN-associated t-SNARE component, has been partially disrupted in *scat* mutant MNs (Figures 4A and S3A). In mammalian cells and yeast, the C-terminal domain of Vps54 interacts with retrograde transport carriers although the precise molecules they bind to on endosomes have not yet been characterized (Perez-Victoria and Bonifacino, 2009; Quenneville et al., 2006). To facilitate the capture of endosome-derived vesicles, the N-terminus of Vps53 and Vps54 interact directly with SNAREs involved in retrograde transport including the t-SNAREs Stx6 and Stx16, as well as the v-SNARE Vamp4 (Perez-Victoria and Bonifacino, 2009). The reduction of GARP components by RNAi in mammalian cells reduces, but does not prevent the formation of TGN-associated t-SNARE complexes (Perez-Victoria and Bonifacino, 2009). We suggest that disruption of *scat* in larval MNs has partially compromised t-SNARE assembly or function contributing to the disruption of LE to TGN retrograde trafficking.

We also demonstrate that some MNs in *scat* loss- and reduction-of-function larvae have defects in the normal localization of the *cis-*Golgi maker Lva, suggesting there is a link between altered retrograde trafficking and Golgi dysfunction (Figure 4D). Similarly, in yeast and humans, Vps51 interacts with Stx6, although disruption of this interaction in yeast does not cause trafficking defects (Conibear et al., 2003; Siniossoglou and Pelham, 2002). Depletion of the zebrafish *Vps51* ortholog, *fat free*, in intestinal cells disrupts vesicle trafficking and Golgi morphology (Ho et al., 2006). The latter suggests that GARP (or EARP) plays an important role in the control of Golgi structure. Analysis of Golgi morphology in the symptomatic wobbler mouse reveals significant vacuolization of the Golgi in the soma of MNs (Palmisano et al., 2011). Finally, in *Drosophila* post-mitotic spermatids, *scat^1^* mutants have defects in the localization of Golgin245, a conserved golgin associated with the TGN, although the *cis-*Golgi marker (the golgin GM130) was unaffected (Fari et al., 2016b). Based on this, we cannot rule out that partial Stx16 mislocalization we observed in *scat^1^* MNs is not due to a more global defect in the integrity of both sides of the Golgi ribbon.

Why does disruption of *scat* expression so strongly impact NMJ development? The membrane trafficking pathway has a well-established function in the control of axon growth and synaptogenesis (Wojnacki and Galli, 2016). Rab5 (EEs), Rab7 (LEs), and Rab11 (REs) all traffic in vertebrate axons and have been implicated in the control of axon growth and guidance (Falk et al., 2014; Ponomareva et al., 2016; van Bergeijk et al., 2015). Membrane and transmembrane proteins are transported to axon terminals via a Rab11-dependent mechanism whereas Rab5 and Rab7 are involved in local recycling and retrograde transport back to the soma (Jin et al., 2018; Khodosh et al., 2006). In *Drosophila*, a loss-of-function mutation in Rab5 causes defects in axon elongation in olfactory projection neurons and sensory neurons (Sakuma et al., 2014; Satoh et al., 2008). Mutations in Rab7 linked to Charcot-Marie-Tooth2b disease cause axon growth and guidance defects in fly sensory neurons (Ponomareva et al., 2016). Rab11 has been implicated in fly models for neurodegenerative disorders including Alzheimer’s Disease (AD) and Huntington’s disease (HD) (Breda et al., 2015; Dumanchin et al., 1999; Greenfield et al., 2002; Li et al., 2012; Li et al., 2009a; Li et al., 2009b; Li et al., 2010; Richards et al., 2011; Steinert et al., 2012). Moreover, *rab11* mutants have defects at the larval NMJ characterized by a significant increase in synaptic bouton number with a clustered phenotype (Khodosh et al., 2006).

We show here that disruption of both *scat* expression and Rab7 function in motor neurons causes a significant reduction in synaptic bouton number (Figure 5A-B) and alters the composition of PSDs at the NMJ (Figures 5C and 6). Boutons are quantified by counting the number of DLG+ synapses at the NMJ, so the reduction in number could be due to loss of DLG from the PSD. That said, we see no evidence of residual HRP+ DLG-axon terminals at these NMJs, suggesting that both the pre- and post-synapse have not developed normally. Interestingly, there is a growing body of evidence suggesting that presynaptic mechanisms are involved in the regulation PSD composition at the fly NMJ. First, presynaptic pMad (*Drosophila* phosphorylated Smad) regulates postsynaptic GluRIIA accumulation in PSDs via a noncanonical BMP signaling pathway (Sulkowski et al., 2016). Second, presynaptic dMon1 regulates levels of GluRIIA at PSDs via a transsynaptic mechanism (Deivasigamani et al., 2015). dMon1 is a conserved effector of Rab5 and is involved in the conversion of EEs to LEs by aiding in the recruitment of Rab7 (Poteryaev et al., 2007; Yousefian et al., 2013). It was proposed that dMon1 may be released from boutons (facilitate the release of some unknown factor) via exosomes in a manner similar to signaling molecules such as Ephrins, Wingless, and Syt4 (Contractor et al., 2002; Korkut and Budnik, 2009; Korkut et al., 2013). Rab7 has also been shown to be required for exosome biogenesis and the release of exosomes, including those that containing miRNAs (Corrigan et al., 2014; Jae et al., 2015). Interestingly, the GARP component *vps-52* regulates miRNA-mediated gene silencing by modulating levels of the miRNA pathway protein GW182 in *C. elegans* (Vasquez-Rifo et al., 2013). This raises the interesting possibility that the synergistic effects we observe in *C380>scat shRNA, Rab7 (DN)* double mutant NMJs may be due to defects in the production and secretion of exosomes containing signaling molecules or miRNAs from presynaptic terminals.

Again, the molecular and cellular mechanisms that lead to MN dysfunction and degeneration in the wobbler mouse remain unknown (Moser et al., 2013; Schmitt-John, 2015). To the best of our knowledge, we have provided the first evidence suggesting that *Vps54* regulates NMJ growth and PSD composition via a Rab7-dependent mechanism. *Vps54* loss-of-function in the mouse causes embryonic lethality with an underdeveloped spinal cord and dorsal root ganglia. These embryos also have severe hypoplasia of the atrial and ventricular myocardium (Schmitt-John et al., 2005). Collectively, these data suggest that Vps54 has a critical function in neuronal and muscular development during embryogenesis. We postulate that the NMJ defects we see in *Drosophila* larvae are analogous to these phenotypes in mice. This observation is not unprecedented in *Drosophila* disease models. There are also synaptic defects that have been observed at the larval NMJ in fly models for ALS, AD, Parkinson’s Disease (PD), and frontotemporal dementia (Shahidullah et al., 2013; West et al., 2015; Zhu et al., 2013). Further investigation into the processes occurring during metamorphosis and in adults is required to determine if loss or reduction of *scat* results in age-progressive MN degeneration.

## AUTHOR CONTRIBUTIONS

Designed experiments: PHP, ECW, ELS, SAB. Performed experiments: PHP, ECW, ELS, MRM. Analyzed the data: PHP, ECW, ELS, SAB. Wrote the paper: PHP, ECW, ELS, JTB, SAB.

## ACKNOWLEDGEMENTS

Some fly strains were from the Bloomington *Drosophila* Stock Center, which is funded by NIH grant P40OD018537. The *scat-RA* cDNA was obtained from the *Drosophila* Genomics Resource Center, which is supported by NIH grant P40OD010949. Some antibodies were supplied by the Developmental Studies Hybridoma Bank, which was created by the NICHD and is maintained by the University of Iowa. The Lva antibody was a gift from John Sisson and Christine Fields (Harvard). Rab5 and Rab11 antibodies were provided by Akira Nakamura (Institute of Molecular Embryology and Genetics). The GluRII antibodies were a gift from Mihaela Serpe (NIH/NICHD). PHP was supported by a Renew DU Postdoctoral Fellowship awarded to SAB and JTB. SAB was funded by a pilot grant from the Knoebel Institute for Healthy Aging at the University of Denver. The authors declare no competing financial interests.

## SUPPLEMENTAL DATA

### SUPPLEMENTAL METHODS

#### Analysis of active zones

To quantify active zone number, NMJs were counterstained with antibodies targeting Brp and HRP as described above. Maximum Z-projections were processed for *scat^1^* mutants at m6/7 in A3 and were quantified by manually counting Bruchpilot (Brp) foci within synaptic boutons. Twenty NMJ’s were analyzed for each genotype in which total Brp punctae and total bouton number per NMJ were counted and analyzed using ImageJ2/Fiji software.

### SUPPLMENTAL FIGURE LEGENDS

**Figure S1.**
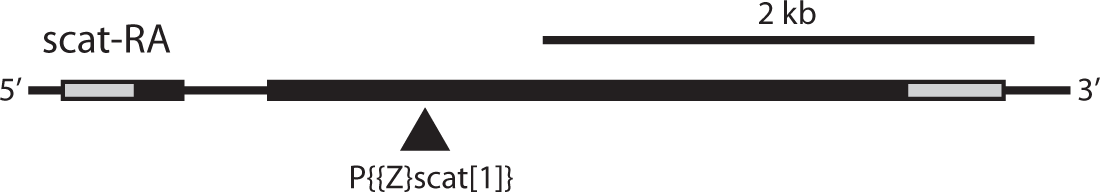
The *scat^1^* allele. This is a diagram showing the location of the P-element insertion in the second exon of the *scat* locus. Using an antibody targeting the N-terminal 200 amino acids of the Scat protein, it has previously been shown that this is a null allele (Fari et al., 2016a).

**Figure S2.**
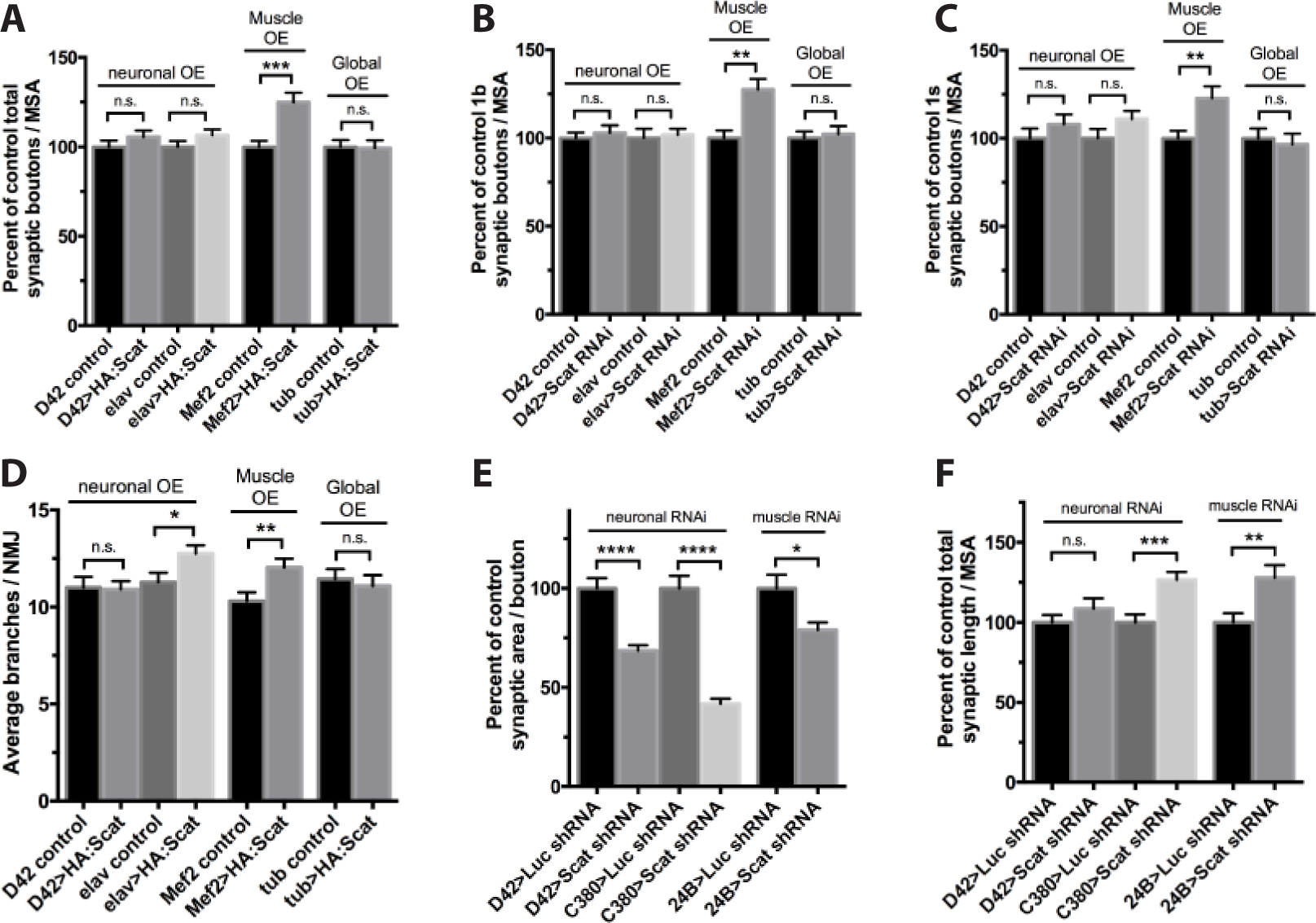
Overexpression of *scat* has an effect on NMJ development. A-F) A *UAS-HA:scat* transgene was expressed using a motor neuron-specific driver (*D42-Gal4*), a strong pan-neuronal driver (*elav-Gal4*), a strong muscle-specific driver (*Mef2-Gal4*), or ubiquitously (*tubulin-Gal4*). The A) total number of boutons, B) type Ib boutons, and C) type 1s boutons are increased when HA:Scat is overexpressed in larval muscle but not when expressed ubiquitously or in neurons. D) The number of branches is significantly increased when HA:Scat is strongly expressed in either neurons or muscle. *N* = 22, 23, 19, 24, 30, 30, 23, and 23 in the order shown in graphs. E) Total synaptic area is increased when strongly expressed in muscle. F) There is no impact on total synaptic length. Synaptic area and length where quantified using the Morphometrics algorithm. *N* = 22, 23, 19, 24, 28, 30, 24, and 27 in the order shown in graphs. Unless otherwise indicated, all statistical comparisons shown have been compared to driver-specific controls (*driver/+* heterozygotes). * p < 0.05, ** p < 0.01, *** p < 0.001.

**Figure S3.**
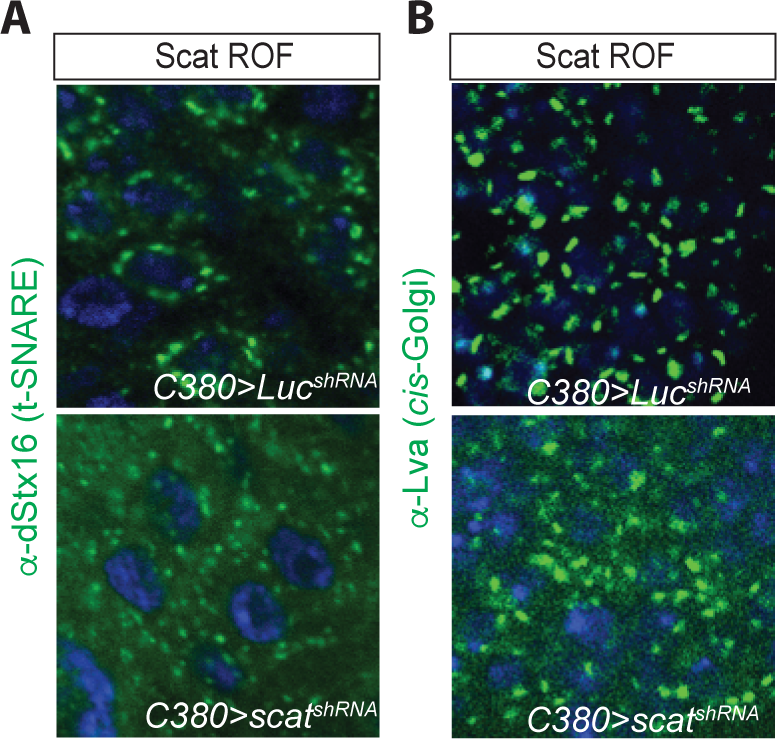
*scat* RNAi disrupts Stx16 localization and Golgi integrity in larval motor neurons. A) Localization of Syntaxin-16 to the TGN is disrupted by motor neuron-specific disruption of *scat* expression using the *C380-Gal4* driver. *C380>Luc^shRNA^* and *C380-scat^shRNA^* lines were stained with an antibody targeting dStx16 (green) and DAPI to visualize nuclei (blue). dStx16 staining is significantly more diffuse (but still clearly punctate) in *scat* RNAi animals compared to controls. B) Localization of Lva is partially disrupted in the cell body of *scat* RNAi motor neurons. Indicated genotypes have been stained with an antibody targeting Lva (green) and DAPI (blue). Scale bar, 2.5 μm.

**Figure S4.**
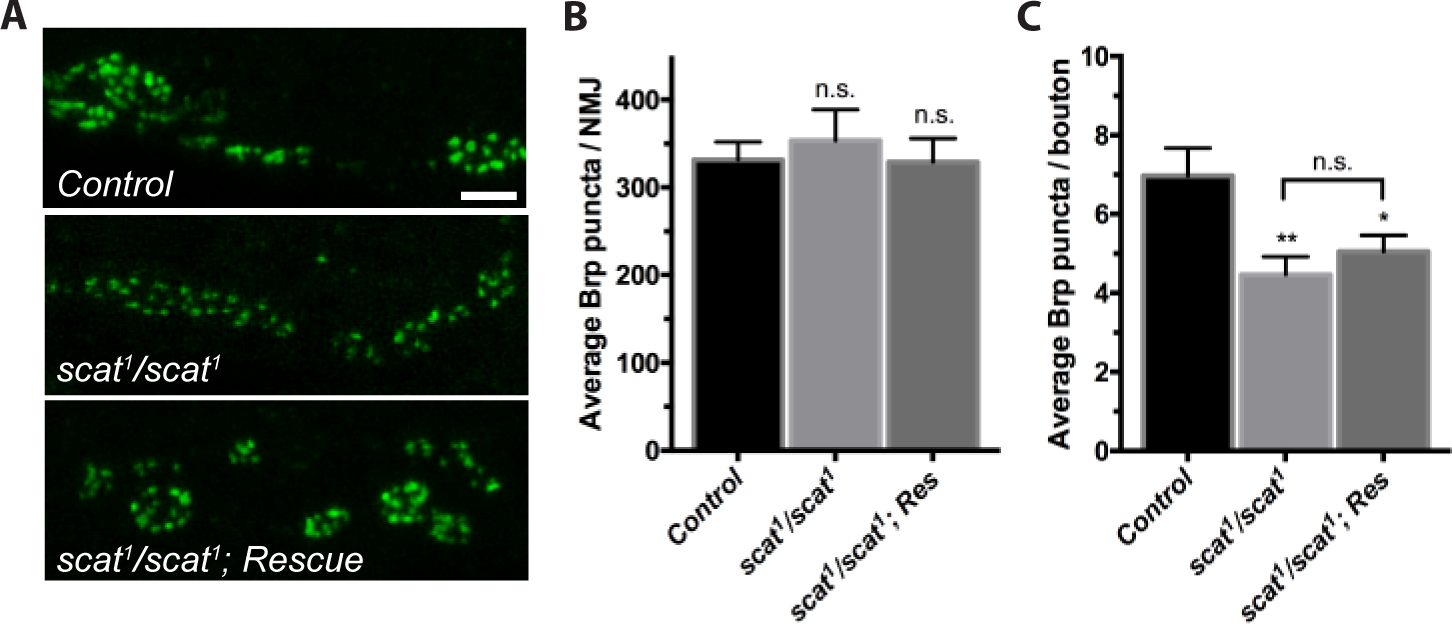
*scat* mutant adults have early-onset locomotor defects and a shorted lifespan. A) NMJs innervating muscle 6/7 in body segment A3 from the indicated genotypes stained with an antibody targeting Brp (green). *scat* mutants have an increased number of small boutons with fewer Brp-positive puncta. Scale bar, 5 μm. B) The total number of Brp-positive punctae per NMJ does not change C) The number of Brp-positive punctae per synaptic bouton increases in *scat* mutants. *N* = 10 for each genotype. Data in graphs are represented by the mean ± SEM. * p < 0.05, ** p < 0.01.

